# Molecular basis for recognition of the Group A Carbohydrate backbone by the PlyC streptococcal bacteriophage endolysin

**DOI:** 10.1101/2021.03.19.436156

**Authors:** Harley King, Sowmya Ajay Castro, Amol Arunrao Pohane, Cynthia M. Scholte, Vincent A. Fischetti, Natalia Korotkova, Daniel C. Nelson, Helge C. Dorfmueller

## Abstract

Endolysins are peptidoglycan (PG) hydrolases that function as part of the bacteriophage (phage) lytic system to release progeny phage at the end of a replication cycle. Notably, endolysins alone can produce lysis without phage infection, which offers an attractive alternative to traditional antibiotics. Endolysins from phage that infect Gram-positive bacterial hosts contain at least one enzymatically active domain (EAD) responsible for hydrolysis of PG bonds and a cell wall binding domain (CBD) that binds a cell wall epitope, such as a surface carbohydrate, providing some degree of specificity for the endolysin. Whilst the EADs typically cluster into conserved mechanistic classes with well-defined active sites, relatively little is known about the nature of the CBDs and only a few binding epitopes for CBDs have been elucidated. The major cell wall components of many streptococci are the polysaccharides that contain the polyrhamnose (pRha) backbone modified with species-specific and serotype-specific glycosyl side chains. In this report, using molecular genetics, microscopy, flow cytometry and lytic activity assays, we demonstrate the interaction of PlyCB, the CBD subunit of the streptococcal PlyC endolysin, with the pRha backbone of the cell wall polysaccharides, Group A Carbohydrate (GAC) and serotype *c*-specific carbohydrate (SCC) expressed by the Group A Streptococcus and *Streptococcus mutans*, respectively.

## Introduction

Endolysins are bacteriophage-encoded PG hydrolases that normally function from within the cell to lyse the bacterial host, releasing progeny phage and completing the phage lifecycle [1]. However, the lytic activity of endolysins can be harnessed for antimicrobial use due to their ability to equally lyse bacteria when applied exogenously, without infection by a parental phage. Due to their direct lytic action on target PG, endolysins are not affected by efflux pumps, alterations in metabolism, or other mechanisms of antibiotic resistance, making them ideal candidates for development against multi-drug resistant organisms [2–4]. Notably, at least three endolysins, some of which are active against methicillin-resistant *Staphylococcus aureus*, are currently being evaluated in human clinical trials for their antimicrobial activity (reviewed in [5]).

Most endolysins, and in particular those from phage that infect Gram-positive bacterial hosts, are comprised of modular domains. An enzymatically active domain (EAD) is generally found in the N-terminal region, while a cell wall-binding domain (CBD) is located in the C-terminal region [6]. As the name implies, the EAD is a catalytic domain that is responsible for cleaving specific bonds in the PG, the nature of which is dependent on the mechanistic class of the EAD. Occasionally, endolysins contain two EADs, although both are not necessarily active. The CBD binds at high affinity [7] to a cell wall-specific epitope and was suggested to dictate genus, species and serovar-specificity of the endolysin. The CBD targets may be surface carbohydrates, wall teichoic acids linked to the Gram-positive bacterial cell wall, or the PG itself [8].

The endolysin now known as PlyC is one of the first described endolysins and remains one of the most studied. In 1934, Alice Evans noted a “nascent lysis” activity derived from streptococcal phage lysates on streptococcal strains that were not sensitive to the phage itself [9]. By 1957, Krause had determined that the phage used by Evans was specific for Group C Streptococci (GCS), but an “enzyme” produced by the phage could lyse Groups A, A-variant, and C Streptococci (GAS, GAVS, and GCS, respectively) [10]. These findings were confirmed by Maxted later the same year and extended to include Group E Streptococci (GES) [11]. In 2001, PlyC, then referred to as the streptococcal C_1_ lysin, became the first endolysin to be used therapeutically, protecting mice from GAS challenge in a nasopharyngeal model [2]. Subsequent studies revealed that PlyC is a structurally unconventional endolysin, which is not encoded by a single gene as found for all other endolysins described to date. Rather, PlyC is a nine-subunit holoenzyme encoded by two distinct genes, *plyCB* and *plyCA*, within a polycistronic operon [12]. Eight PlyCB subunits self-assemble into a ring structure and form the basis of the CBD that binds the streptococcal surface. A single PlyCA subunit contains two distinct EADs separated by an extended α-helical linker region, which interfaces with the N-terminal residues of the PlyCB octamer [13]. The PlyCA EADs consist of a <u>g</u>lycosyl <u>h</u>ydrolase (GH) domain and a <u>c</u>ysteine, <u>h</u>istidine-dependent <u>a</u>minohydrolase/<u>p</u>eptidase (CHAP) domain. The very high lytic activity of PlyC relative to other endolysins is attributed to synergy/cooperativity between the two EADs.

Although the EADs of PlyCA have been extensively characterized with respect to specificity of PG bonds they cleave, active-site residues, and their synergistic activity, it is unclear how PlyCB recognizes PG of GAS, GCS and GES. Similar to other Gram-positive bacteria, the plasma membrane of streptococci is surrounded by thick cell wall that consists of a complex network of PG with covalently attached polysaccharides. Rebecca Lancefield utilized the unique immunogenicity of the surface carbohydrates in β-hemolytic streptococci to subsequently separate them into serogroups [14]. *S. pyogenes* is categorized as Group A Streptococcus, whilst *S. dysgalactiae subsp equisimilis* (SDSE) produce at least two types of carbohydrates and are annotated as Group C and G Streptococci, respectively [15]. Pioneering work by many researchers have revealed that the cell wall polysaccharides of GAS, GCS and *S. mutans*, consist of a pRha backbone modified with species-specific and serotype-specific glycosyl side chains (Fig. 1A) [16, 17]. In GAS, GCS and *S. mutans* serotype *c*, the polysaccharides termed the Lancefield group A carbohydrate (GAC), group C carbohydrate (GCC), and SCC, respectively, have a conserved repeating →3)α-Rha(1→2)α-Rha(1→ di-saccharide backbone [16, 17]. The β-*N*-acetylglucosamine (GlcNAc) side chains are attached to the 3-position of the α-1,2 linked rhamnose (Rha) in GAC [16, 18, 19]. The GCC side chains have two *N*-acetyl-galactosamine (GalNAc) residues attached to the same hydroxyl of Rha [16, 18, 20]. SCC carries the α-glucose (Glc) side chains attached to the 2-position of the α-1,3 linked Rha in SCC [17]. Additionally, the side chains of GAC and SCC are decorated in parts with the glycerol phosphate (GroP) moiety [21]. The GAC and SCC biosynthetic pathways are encoded by the 12-gene loci *gacABCDEFGHIJKL* and *sccABCDEFGHMNPQ*, respectively. The molecular mechanisms of GAC and SCC biosynthesis have been the focus in a number of recent studies [21–25].

**Fig 1).**
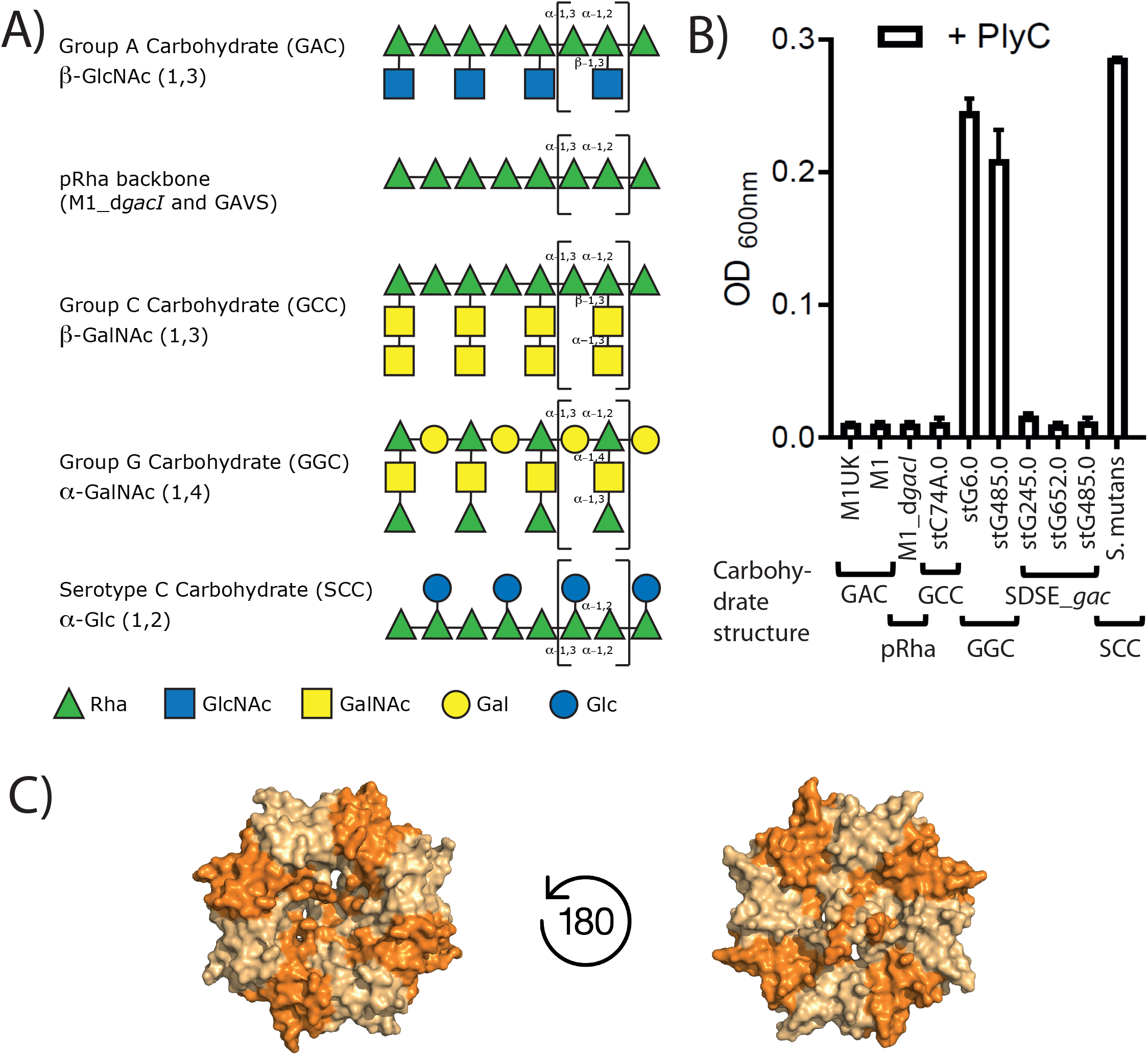
A) Symbolic drawings of the carbohydrate structures of GAS (GAC), polyrhamnose backbone (pRha) in GAVS and M1_d*gacI*, GCS (GCC), GGS (GGC) and *S. mutans* (SCC). For simplicity, the reported glycerol phosphate of occasionally present on the GAC and SCC side chains have been omitted. Repeat units are marked with brackets. The pRha backbone with alternating (α1->2) and (α1->3) linkages, whilst the α1->2 Rha being decorated with a 11->3 side chain is a commonality among all PlyC susceptible strains. B) PlyC lysis of streptococcal pathogens. Group A and Group C Streptococci serotypes are susceptible to PlyC-mediated lysis. Group G Streptococci show limited susceptibility and *S. mutans* is resistant to PlyC lysis. Three SDSE isolates that produce the GAC instead of GGC (SDSE_*gac*) are susceptible to PlyC treatment. C) Bottom and top view onto WT PlyCB octameric structure with the GlyH and EAD domains omitted (4f78.pdb).

Remarkably, extracted GAC is known to partially inhibit the lytic actions of PlyC [26]. Furthermore, a GAS mutant carrying the unmodified GAC (lacking the GlcNAc side chains) displayed enhanced susceptibility to PlyC [25], implicating PlyCB in recognition of the pRha backbone. Here, we identify the pRha backbone of GAC as the definite minimal binding epitope for PlyCB.

## Materials and Methods

### Bacterial strains, growth conditions and media

*Streptococcus pyogenes* strain D471 was propagated on solid media in plates containing Todd-Hewitt broth supplemented with yeast extract (0.2%) (THY) and agar (1.4%) or in liquid THY as described by Gera et al [27]. *S. mutans* wild type (Xc) and mutants were grown in Todd-Hewitt broth with 1% yeast extract. All cultures were grown without antibiotics and without aeration at 37°C. *E. coli* genotypes DH5*α* (NEB, cat. No. C2988J), DH10*α* (Thermo Fisher, cat No. 18297010), BL21 (NEB, cat No. C2530H) and Origami 2 (Millipore Sigma, cat. No. 71346) were used for routine plasmid propagation or protein expression and grown in Lysogeny Broth (LB) medium supplemented with either 50 µg/ml kanamycin, 100 µg/ml ampicillin or 35 µg/ml chloramphenicol as needed.

Bacterial strains *E. coli* CS2775 were transformed with pRGP1 plasmid [28], *gacABCDEFG* [23] to produce pRha or empty plasmid control (pHD0131). The bacterial cells were grown overnight in LB containing erythromycin (150 µg/ml) at 37°C and used next day for whole cell Western blots and FACS and microscopy analysis.

### Recombinant expression and purification of PlyCB_WT_ and PlyCB_R66E_

PlyCB_WT_ (GenBank ID: NC_004814.1:7517-7735) was expressed from a pBAD24 vector and purified from BL21 cells as previously described [12]. In brief, the culture was grown in LB and induced with 0.25% L-arabinose at OD_600_ ~1.2-1.4 (Alfa Aesar, cat no. A11921). Cultures were grown at 37°C with shaking at 180 RPM for 3-4 hours, centrifuged at 4500 x g and resuspended in phosphate-buffered saline (PBS). Lysis was performed by a French press (1800 psi). Benzonase (Millipore Sigma, cat. No. 70746-3) was added and the lysate incubated at room temperature with rotation for 20-30 min. The lysate was centrifuged at 20,000 x g for 20 min and the cleared lysate was passed through a 0.45-µm filter and loaded onto a XK-26/20 column (Cytiva) with 30-35 ml ceramic hydroxyapatite (Bio-Rad, cat no. 1582000). PlyCB_WT_ was eluted from the column with three column volumes of 1 M sodium phosphate buffer (pH 7.2). Protein was subsequently dialyzed in PBS, 10% glycerol and stored at −80°C until use. Protein purification of PlyCB_R66E_ was performed as PlyCB_WT_ [13]. The fluorescent labeling of PlyCB_WT_ and PlyCB_R66E_ was performed using the manufacturer’s recommended guidelines (Thermo Fisher, cat. No. A20174). Analytical gel filtration was used to determine the multimeric nature of PlyCB and PlyCB_R66E_. A total of 100 µl (0.1 mg) of each protein was applied to a Superose 12 10/300 GL column (Cytiva) and run in isocratic conditions in PBS buffer for 1.5 column volumes on an AKTA FPLC system (Cytiva). Gel filtration standards (Bio-Rad) were also run under identical conditions.

### Purification of GAC

GAS cells were grown overnight in THY media at 37°C. Cultures were centrifuged at 4,500 x g. Pellets were washed and resuspended in 40 ml distilled water per each original liter of media used and combined in an 800 ml beaker. 22.5 ml 4N sodium nitrite (5 ml per liter of culture) was added to the beaker in addition to 22.5 ml glacial acetic acid (5 ml per liter of culture). An orbital shaker was used to gently mix the beaker for 15 min in a hood. The mixture was centrifuged in 500 ml bottles at 8000 x g for 15 min. The supernatant was decanted to a new beaker and neutralized with 1M sodium hydroxide. The total solution, about 300 ml, was filtered with a 0.45-micron filter assembly. 50-50 ml aliquots were deposited in a 3.5 kDa membrane and dialyzed in a 4-liter beaker overnight with water. The following day, the solution was concentrated using an Amicon 400 ml stirred cell (model 8400) filter assembly with a 76 mm diameter Ultracel^®^ 5 kDa ultrafiltration disc (Millipore Sigma, cat. No. PLCC07610) for 2 hours with 60 psi. The ~10 ml concentrate was loaded onto an S-100 column for final purification. Fractions were assayed for Rha, lyophilized and stored at 4°C until use.

### Calculation of pRha concentration

A modified protocol by Edgar *et al*. [21] based on a protocol from DuBois *et al*. [29] was used to determine Rha concentration in purified GAC. Briefly, anthrone reagent was prepared by dissolving 0.2% w/w anthrone in H_2_SO_4_. Eighty microliters of aqueous samples or standards containing either GAC or L-Rha at known concentrations were added to a 1.5 ml microfuge tube. To this same tube, 320 µL of the anthrone reagent was added. Samples were boiled at 98°C for 10 min in a heat block. Samples were cooled to room temperature, transferred to a quartz plate, and the absorbance at 580 nm was recorded using a spectrophotometer. Rha concentration was interpolated using an L-Rha standard curve.

### Precipitation of PlyCB with GAC

PlyCB_WT_ and PlyCB_R66E_ samples were defrosted from storage at −80°C. Lyophilized GAC was resuspended in PBS. Both proteins and GAC were added to a 3.5 kDa dialysis membrane and dialyzed overnight in PBS. Protein concentrations were determined using a NanoDrop spectrophotometer (Thermo Fisher ND-2000) at 280 nm and were diluted to 5 mg/ml. The GAC was also assayed and diluted with PBS to 1.6 mg/ml. One-hundred microliters of proteins and 100 µl of GAC or PBS were mixed in a 250 µl quartz plate and allowed to incubate without shaking at room temperature. Visible precipitate formed in samples in 5-8 minutes. After recording the precipitate at 340 nm using a Spectramax^®^ M5 (Molecular Devices) spectrophotometer, the total sample volume was transferred to a 1.5 ml microfuge tube. Samples were centrifuged at 14,000 x g to pellet the precipitate and supernatants were transferred to new 1.5 ml microfuge tubes. Two-hundred microliters of 8 M urea was added to the pellet. Pellets were resuspended and 5 µl of either pellet or supernatant were added to 40 µl water with 8 µl 6x Laemmli buffer with DTT. Samples were boiled at 98°C for 8 min, and then 12.5 µl were loaded onto a 7.5% SDS-PAGE and run for 32 min at 200 V. Proteins were visualized using Coomassie stain.

### Lysis assay

A turbidity reduction assay was used to ascertain strain sensitivity to PlyC. This assay was performed as previously described [30], except PlyC was used at 2 µM. Eight technical replicates were performed.

Sensitivity of streptococcal species to PlyC-mediated lysis was analyzed using a wide range of clinical isolates: 1) GAS isolates: M1_UK_, WT 5448 strain [M1], delta*gacI* 5448 strain [d*gacI*]; 2) GCS isolate: stC74A.0; 3) SDSE_*gac* isolates: stG245.0, stG652.0. stG485.0; GGS: stG6.0, stG485.0 and 4) *S. mutans* serotype c were used as negative controls. Briefly, all streptococcal strains were grown in THY at 37°C overnight in 5% CO_2_, except for *S. mutans*, which was grown in THB media. Next day, the bacterial cells were inoculated in 1:100 fresh media and grown until mid-logarithmic phase (OD_600_ 1.0). The cells were washed in PBS and resuspended to an OD_600_ of 2.0. In a 96-well plate, to a 100 µl of bacterial cells, 100 µl of PlyC [1 µg/ml] was added and immediately read at an absorbance of OD_600_. The obtained values were standardized by subtracting from the background values. The data is plotted using GraphPad Prism version 9.

### SDS-PAGE and blotting analysis

PlyCB_WT_ binding to recombinant *E. coli* expressing pRha was conducted using blot analysis. Briefly, the lysate from the overnight cultures was analyzed in 20% tricine gels. SDS-PAGE and protein transfers were performed according to manufactures instructions, Atto Ae-6050 Mini Gel chamber and Novex protein separation from Thermo Fisher, respectively. The PVDF membranes were blocked with 5% non-fat dry milk with Tris-Buffered Saline, 0.1% Tween^®^ 20 detergent prior to incubation with PlyCB_WT_ labelled with Alexa Fluor^®^ 647 (1:1000) for one hour at room temperature. Goat anti-rabbit GAC antibodies conjugated with IRDye^®^ 800CW were used as a positive control (abcam ab216773). The resulting blots were imaged using the Licor Odyssey FC Imaging System. All the blots were processed in parallel under the same conditions.

### Microscopy

Microscopic analysis of *E. coli* bacteria was performed using cells that were grown overnight in LB supplemented with antibiotics at 37°C and diluted 1:100 the next day and regrown until OD_600_ reached 0.5. The cells were washed twice with PBS for 5 min at 10,000 rpm and stained with 1:1000 dilution of PlyCB^AF647^ and left for 20 minutes on ice in the darkness. Prior to fixing the cells with 4% paraformaldehyde, the cells were washed again with PBS. The fixed cells were mounted on 1% agarose coated microscopic slides and viewed under the CY5 channel on a fluorescent Deltavision widefield microscope.

### FACS analysis

*E. coli* cells were grown overnight as described above and diluted 1:100 the next day, grown at 37°C and used for the assay at OD_600_ = 0.5. The cells were washed with PBS and probed with PlyCB^AF647^ or PlyCB_R66E_^AF647^ at 1:1000 dilution at 1:1000 dilution. Anti-GAC antibodies conjugated with FITC (ABIN238144, antibodies-online, titer 1:50) were used as a positive control. The samples were incubated for 20 minutes on ice at dark conditions. The cells were washed twice with PBS at 14,000 rpm for 5 minutes and fixed with 4% paraformaldehyde. BD LSRFortessa Flow Cytometry software was used to analyse the samples and the data interpretation was conducted with FlowJo™ software v10.6.2.

### PlyC hydrolysis of sacculi

*S. mutans* wild-type (WT) and the mutant strain sacculi were obtained by the SDS-boiling procedure [25] followed by four washes each with 1 M NaCl and distilled water. The sacculi were resuspended in PBS to OD_600_ of 1.0 and incubated with PlyC (5 µg/ml) in a 96-well plate. The lysis was monitored after 10, 20, 30, 40, 50 and 60 min as a decrease in OD_600_. Results were reported as fold change in OD_600_ of the sacculi incubated with PlyC vs the sacculi incubated without PlyC.

## Results & Discussion

### Pathogenic streptococci producing the GAC are susceptible to PlyC

A major component of the GAS cell wall is the GAC, building approximately 50% of the cell wall by weight [31]. The GAC is universally conserved amongst all isolated GAS strains on the basis of the gene cluster sequence conservation [32]. A number of *S. dysgalactiae subsp. equisimilis* (SDSE) isolates, naturally belonging to Group G Streptococci (GGS), have been reported to have undergone homologous recombination and replaced their Group G Carbohydrate (GGC) gene cluster in parts with the GAC gene cluster [33–35]. We therefore expanded the previously reported PlyC streptococci cell lysis assay used by Nelson *et al*. [2] to investigate those new isolates named SDSE_*gac*. We also tested if PlyC was able to lyse a selection of GAS serotypes including a newly emerged isolate M1_UK_ [36], and included negative controls GGS isolates and *S. mutans* serotype C (Fig. 1B). In agreement with the published literature, all tested GAS serotypes are susceptible to PlyC lysis and the two GGS isolates are resistant to PlyC. The GGC does not contain the GAC, GCC and SCC pRha backbone with →3)α-Rha(1→2)α-Rha(1→ di-saccharide repeats, but an alternating Rha-Gal carbohydrate [37] (Fig. 1A). All three SDSE_*gac* isolates tested have inherited parts of the *gac* gene cluster and produce the GAC instead of the GGC [34]. Strikingly, the SDSE_*gac* strains are all sensitive to PlyC treatment. The fact that PlyC is able to lyse SDSE_*gac* strains that express the GAC, but PlyC does not lyse the isogenic GGS strains producing the GGC strongly suggests that the GAC is a critical component of PlyC recognition and subsequent activity.

Importantly, all strains tested in this study that are susceptible to PlyC lysis produce a cell wall polysaccharide that contains the pRha backbone and a β-linked sugar substituent on the *α*1,2-linked Rha (Fig 1A). We therefore suggest that the pRha backbone with and without a side chain are both vital ligands to assist PlyC activity and the new SDSE_*gac* isolates will also be susceptible to PlyC treatment due to production of the GAC.

### Purified GAC precipitates PlyCB - but not PlyCB_R66E_

The lysis assay of GAS cells, and in particular of the SDSE_*gac* variants, suggests that either the ubiquitous pRha or GAC in GAS cells is the ligand for PlyCB. We propose that the PlyCB octameric CBD binds GAC and/or the GAC pRha backbone. We tested this hypothesis by investigating the binding of PlyC to partially purified GAC. We hypothesized if the GAC was able to precipitate PlyCB, an interaction of the two systems must have occurred [38]. As a negative control, we employed the previously published inactive mutant PlyCB_R66E_, which lost the ability to bind to GAS cells [13]. Consistent with prior analytical gel filtration and dynamic light scattering results [12], as well as the crystal structures [13, 39, 40], both PlyCB and its R66E mutant self-assemble into stable octameric structures (Fig 1C, SF 1). The purified proteins were incubated with the extracted GAC, and precipitation was monitored at 340 nm, a standard wavelength for measuring protein aggregation [41, 42] (Fig. 2A, B). Whilst keeping the PlyCB concentration constant, we varied the concentration of GAC. Within five minutes at room temperature the solution became turbid, suggesting aggregation (Fig. 2A). When the PlyCB concentration was kept constant and the GAC concentration was varied, the turbidity correlated with PlyCB concentration in a dose dependent manner, suggesting that PlyCB requires GAC to aggregate. Importantly, PlyCB did not self-aggregate when no GAC was added in the assay. Furthermore, no aggregation was detected when PlyCB_R66E_ was incubated with purified GAC (Fig. 2B). To demonstrate the presence of PlyCB in the precipitates, we analyzed the soluble and pellet fractions (Fig. 2C, D). A higher yield of aggregated PlyCB was found in the pelleted samples when compared to the soluble fraction (Fig. 2D). A similar precipitation effect was observed when we varied the PlyCB concentration and kept the GAC concentration constant (Fig. 2E, F), demonstrating that both species are necessary for an interaction.

**Fig 2).**
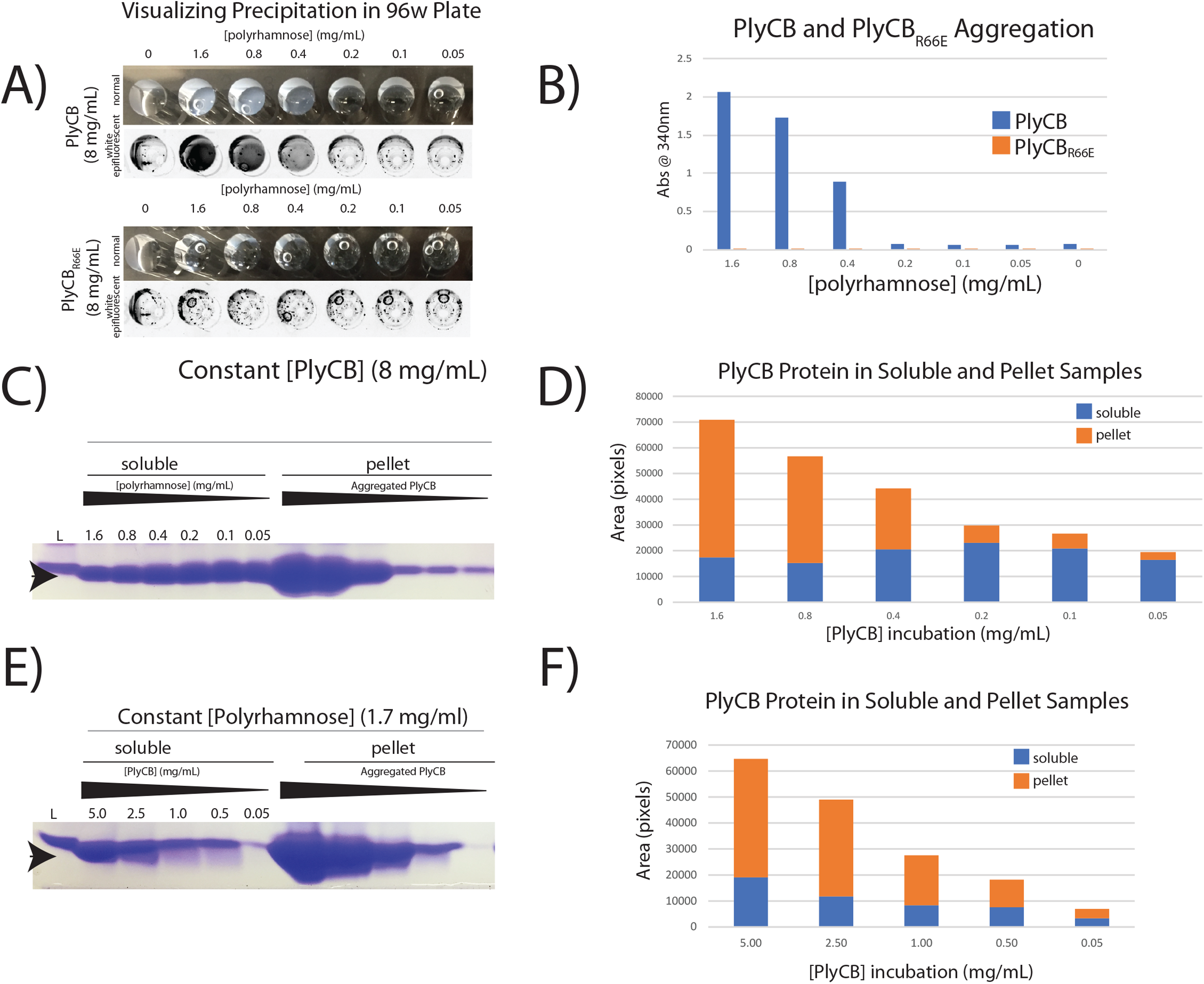
Precipitation studies of purified PlyCB and GAC reveal direct interaction of PlyCB with GAC. A) The PlyCB concentration is kept constant whilst the GAC concentration is varied. Visible precipitate forms at the higher concentrations. B) The precipitate level is measured spectrophotometrically at 340 nm and compared to the mutant PlyCB_R66E_, which does not bind the GAC. C) Coomassie stained and D) densitometry analysis of PlyCB protein from the supernatant fraction and aggregates (pellets). E, F) The same dose dependency is observed when the PlyCB concentration is varied. Arrowhead depicts PlyCB protein at 8 kDa.

### PlyCB binds to recombinantly produced pRha backbone

The purified GAC from bacteria contains a mixture of carbohydrates, including the fully decorated GAC with GroP[21] and a small proportion of the polysaccharide lacking the side chains [43]. PlyC is able to lyse a number of GAS mutants including GAVS and d*gacI*_M1 [2, 11, 25, 44], that decorate the cell wall with the unmodified GAC lacking the side chains (Fig. 1A, B), suggesting that the pRha backbone of the GAC is the minimal carbohydrate structure required for PlyCB binding. To test this hypothesis, we recombinantly produced the pRha backbone in *E. coli* cells. We and others have previously reported that the *S. mutans sccABCDEFG* gene cluster, when transformed into *E. coli* cells, functionally produces the pRha backbone attached to the lipid A [23, 28]. Additionally, to understand if PlyCB recognizes a specific pRha backbone, we engineered *E. coli* cells expressing the GAC gene cluster *gacABCDEFG* required for the GAC pRha production. *E. coli* cells carrying an empty plasmid were used as a negative control.

Next, we investigated the binding of PlyCB conjugated with Alexa Fluor^®^ 647 (PlyCB^AF647^) to an *E. coli* total cell lysate expressing the pRha backbone of the SCC or GAC, respectively (Fig. 3A). The blotted membranes were incubated with PlyCB^AF647^, and a positive interaction between PlyCB^AF647^ and the *E. coli* produced pRha is visualized as a prominent band around 40 kDa. The size of the band agrees with the band detected by anti-GAC antibodies that were previously reported to recognize pRha [23] (Fig. 3B). Importantly, PlyCB^AF647^ and GAC antibodies do not interact with the cell lysate of *E. coli* expressing an empty plasmid (Fig. 3 A, B). Contrary, the R66E-mutant protein was not able to detect the pRha of the PAGE separated sample (SF 2). We further confirmed the ability of PlyCB to bind to *E. coli* cells decorated with the pRha by fluorescent microscopy (SF 3). Only cells that produce the pRha are detected by the PlyCB^AF647^, in agreement with the results of the blot analysis.

**Fig 3).**
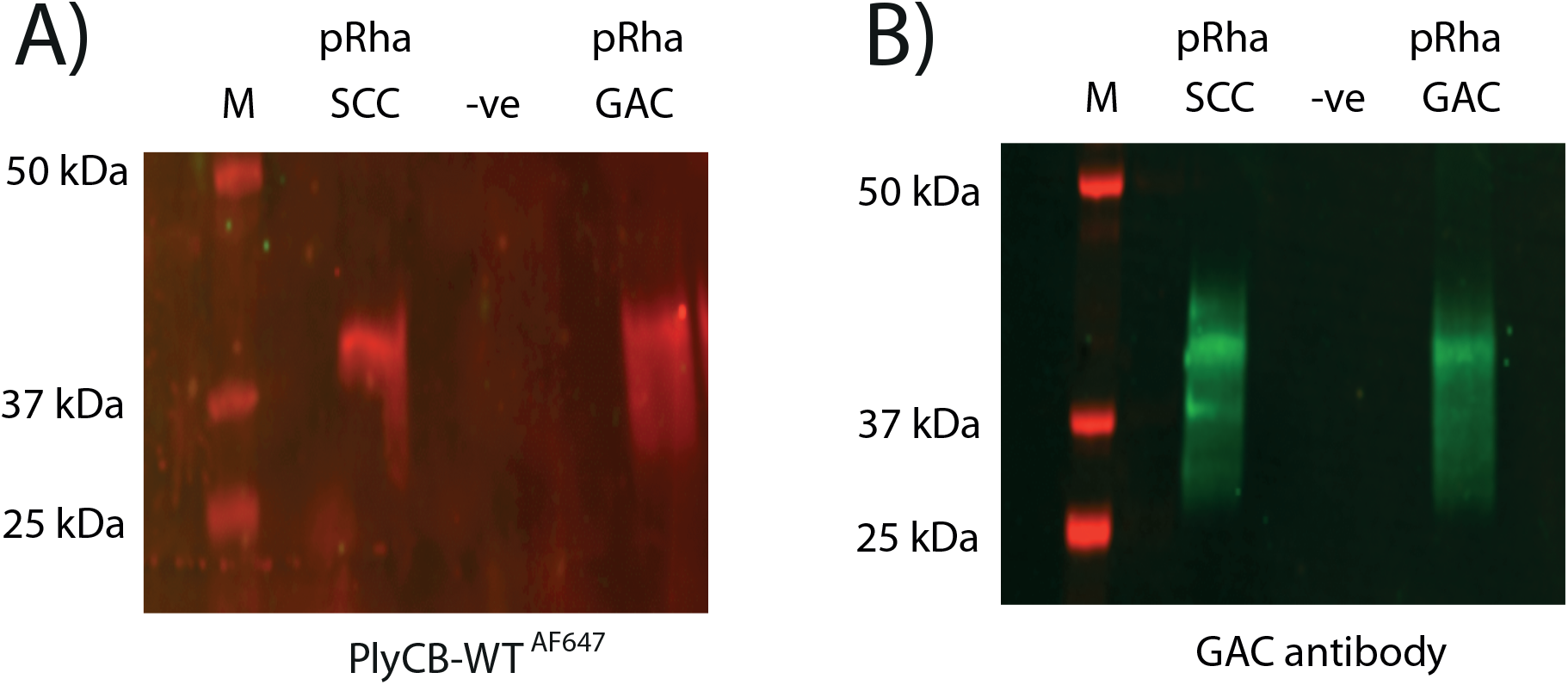
Representation of immunoblot analysis of the cell lysate of *E. coli* expressing the SCC and GAC pRha and carrying an empty control plasmid (−ve). A) Blot was incubated with PlyCB-Alexa Fluor^®^ 647 (PlyCB_WT_^AF647^). B) Probing the same samples with the GAC antibodies confirms the presence of GAC in the bands. Molecular mass markers are given in kDa.

To gather additional evidence that PlyCB interacts with the pRha backbone, we established a flow cytometry assay to analyse the binding of PlyCB^AF647^ to pRha-producing *E. coli*. Unstained *E. coli* cells that express pRha or carry an empty plasmid sort in the identical range (Fig. 4A). The GAC antibodies label exclusively the cells producing pRha (Fig. 4A). A similar pattern of the GAC antibodies binding is observed when the cells were incubated with PlyCB^AF647^ (Fig. 4B). Contrary, the PlyCB_R66E_^AF647^ mutant protein does not bind to *E. coli* cells, and PlyCB^AF647^ does not interact with the cells expressing an empty vector (Fig. 4B). Taken together, these data provide the first definitive evidence that the pRha backbone of GAC and SCC is a binding receptor of the PlyCB octameric subunit.

**Fig 4).**
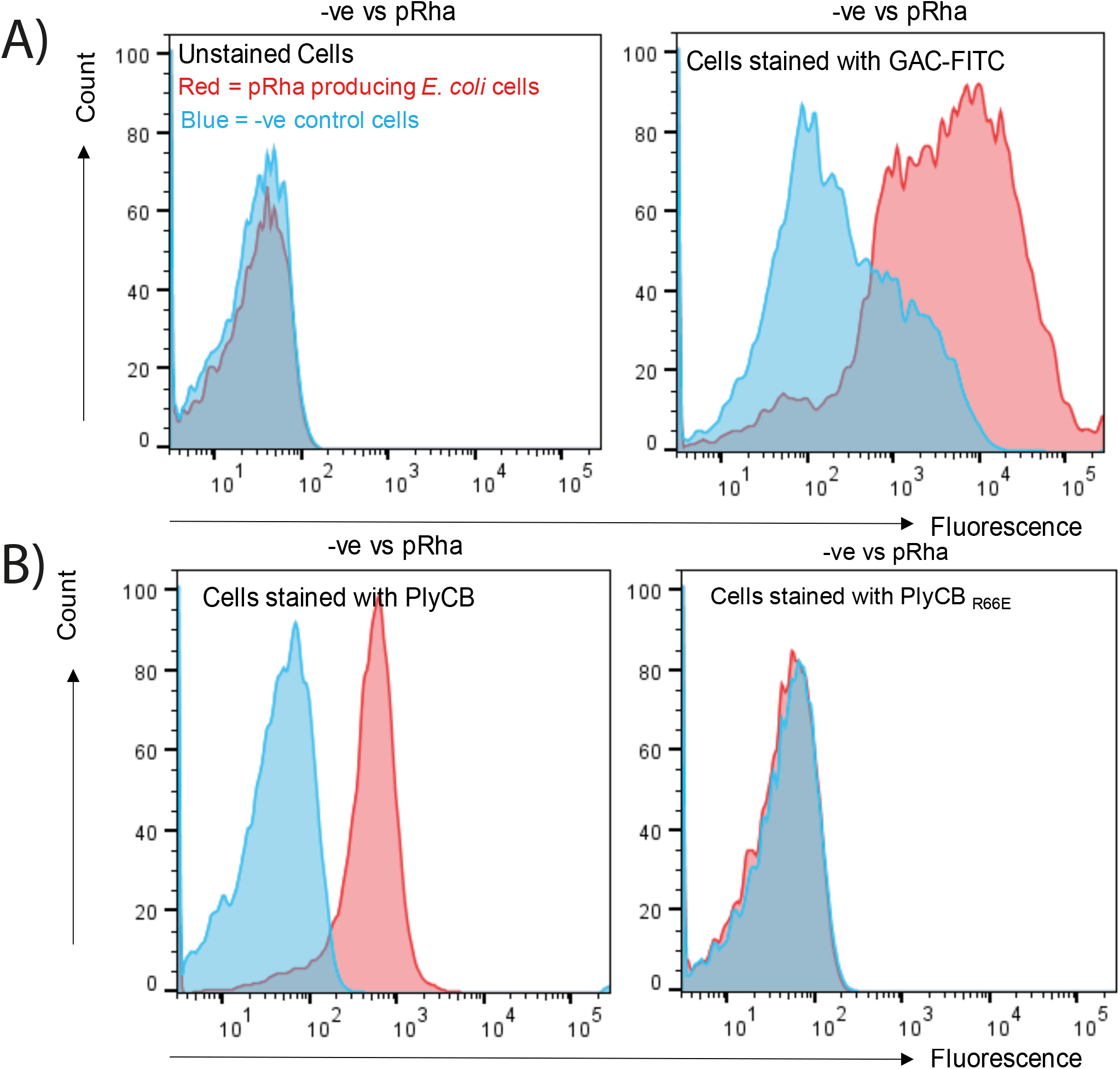
PlyCB binding to *E. coli* cells were investigated by flow cytometry after labelling with PlyCB_WT_^AF647^ and PlyCB_R66E_^AF647^ mutant proteins. Blue: −ve control cells without pRha. Red: pRha producing *E. coli* cells. Representative histograms are shown. A) Left panel: unstained cells. Right panel: The anti-GAC antibodies (GAC-FITC) were used as a positive control to label the *E. coli* cells producing pRha. The antibodies do not bind to the *E. coli* cells carrying an empty plasmid (−ve).B) Left panel: PlyCB_WT_^AF647^ binds to the *E. coli* cells producing pRha, but not to the *E. coli* cells carrying an empty plasmid (−ve). Right panel: PlyCB_R66E_^AF647^ does not binds to the *E. coli* cells producing pRha.

### PlyC lyses engineered *S. mutans* producing the GAC

Despite the fact that the SCC pRha backbone is identical to the GAC, *S. mutans* is resistant to PlyC lysis (Fig. 1A). To get a better understanding why *S. mutans* is resistant to PlyC, we compared PlyC-induced lysis of the sacculi purified from *S. mutans* WT and a number of mutant strains producing different SCC variants (Fig. 5). First, we examined the Δ*sccH* mutant producing the GroP-deficient SCC [21]. Similar to *S. mutans* WT, Δ*sccH* was resistant to PlyC-mediated lysis (Fig. 5). Second, we tested the *sccN* deletion mutant, Δ*sccN*, that is deficient in the enzyme required for generation of the Glc side chains [45]. A time dependent lysis is observed for Δ*sccN*. Expression of the WT copy of *sccN* in Δ*sccN* (the Δ*sccN*:p*sccN* strain) fully restored the resistance of the bacteria to PlyC (Fig. 5). These observations clearly suggest that PlyC is able to bind to the *S. mutans* cells producing the unmodified pRha backbone, and the Glc side chains in SCC hinders PlyC binding. We then investigated whether the addition of the GAC GlcNAc side chains to the pRha backbone affects sensitivity of the engineered *S. mutans* sacculi to PlyC-induced lysis. We expressed the GAS genes *gacHIJKL* required for the formation and addition of the GlcNAc side chains and GroP to GAC [21], in the Δ*sccN* background strain in two versions, creating the Δ*sccN:pgacHI*JKL* and Δ*sccN:pgacHIJKL* strains [45]. The plasmid *pgacHI*JKL* contains an inserted stop codon in the *gacI* gene required for generation of the GlcNAc side chain, and, therefore, the Δ*sccN:pgacHI*JKL* strain produces the unmodified SCC lacking any side chains (Fig. 5). As expected, the sacculi isolated from this strain remains susceptible to PlyC lysis. We previously showed that in Δ*sccN:pgacHIJKL*, the Glc side chains are replaced with the GlcNAc side chains [45]. Interestingly, expression of *gacHIJKL* in Δ*sccN* did not restore the resistance of the bacteria to PlyC (Fig. 5), indicating that the GlcNAc side chains do not obstruct PlyC binding. Lastly, we analyzed PlyC-mediated lysis of sacculi purified from the Δ*rgpG* mutant, which is deficient in SCC expression [45]. The RgpG protein catalyzes the first step in SCC biosynthesis [46]. In comparison to Δ*sccN*, PlyC-induced lysis of Δ*rgpG* was less pronounced (Fig. 5), indicating the importance of the pRha backbone of SCC in PlyC activity and supporting the findings that the pRha backbone is a ligand contributing to PlyC binding. These studies reveal that if the SCC is ‘unmasked’ *i*.*e*., stripped of the Glc and Glc-GroP side chains, it becomes a ligand for PlyCB and that *S. mutans* is PlyC susceptible if SCC is replaced with GAC.

**Fig 5).**
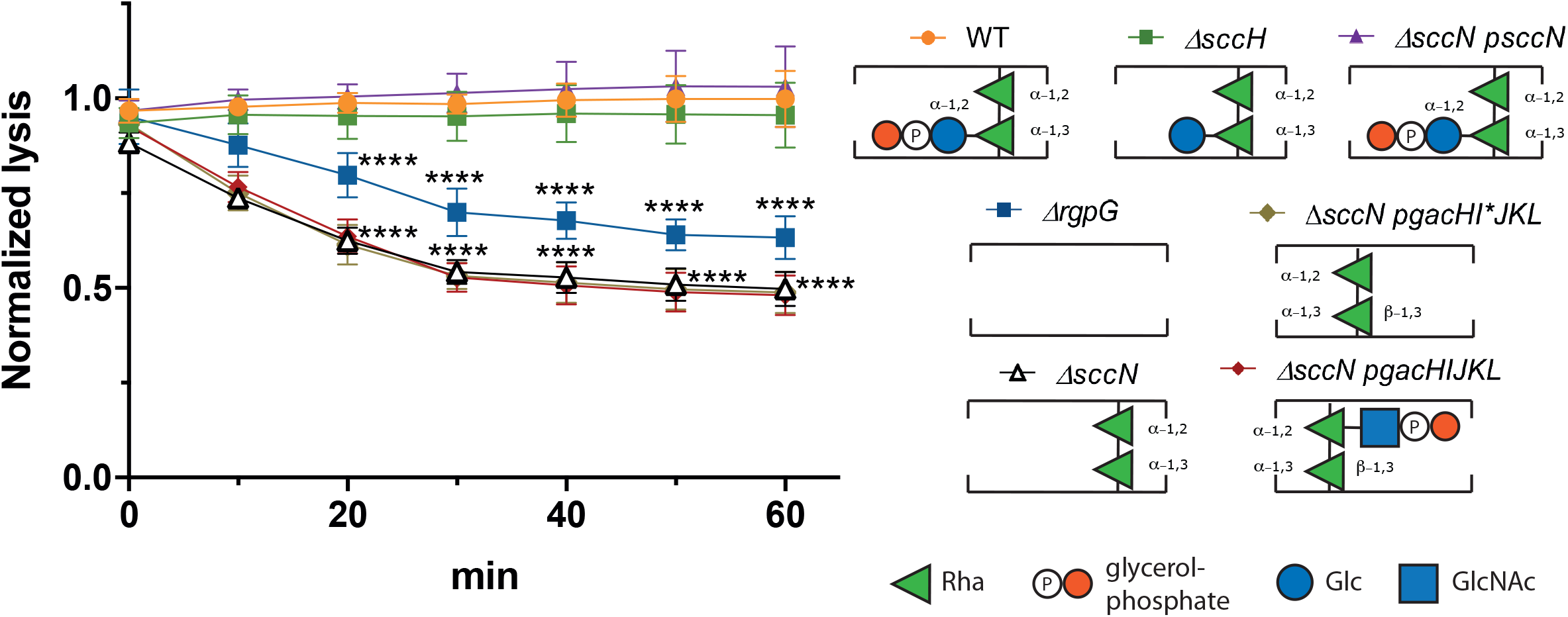
The PlyC-mediated lysis of sacculi purified from *S. mutans* strains. The lysis was monitored after 10, 20, 30, 40, 50 and 60 min as a decrease in OD_600_. Results are presented as a fold change in OD_600_ of the sacculi incubated with PlyC *vs*. the sacculi incubated without PlyC. Data points and error bars represent mean values of four biological replicates and standard deviation, respectively. P-values were determined by two-way ANOVA with Dunnett’s multiple comparisons test.

### Concluding remarks

A structural feature of the PlyCB protein remains to be discovered that explains why only certain streptococci are susceptible to PlyC’s lytic activity. The pRha decorated with an *α*–linked side chain sugar appears not compatible with the PlyCB ligand binding site and therefore only those streptococci expressing pRha decorated with β-linked substituents, such as GlcNAc and GlcNAc-GroP are susceptible. This could potentially be exploited for diagnostic purposes or in the case of SCC and Group G Streptococci, opens up the potential for novel therapeutic approaches. If SCC was treated by PlyC in combination with an additional enzyme that removes the *α*–linked side chains in these streptococcal carbohydrates, this would expose the pRha backbone and subsequently make these strains susceptible. A recently accepted manuscript by Boendum *et al*. [40] reported 19 potential binding states of tetrarhamnose to PlyCB. Once the pRha ligand site has been identified, this could also be further exploited by directed evolution approaches to generate PlyCB protein variants that are capable to bind the carbohydrates from, for example, SCC and Group G Streptococci.

Whilst much has been learned about the structure and function of PlyC in the past 20 years, many questions remain, specifically with respect to its interaction with the PG. Considering the average length of the cellular pRha is 7-10 kDa [25] and that the *α*1,2-1,3-pRha with or without β-configured GlcNAc/GalNAc-side chains bind to PlyCB, it is inviting to speculate that an element of avidity is responsible for tight binding of the PlyCB octamer to the streptococcal surface, in agreement with the recent accepted manuscript [40]. Further proof is needed to substantiate this hypothesis. Another question lies in the actions of the EADs relative to the PlyCB octamer. PlyC clearly has a high turnover as demonstrated in multiple biochemical assays. However, it is unknown if PlyCB “docks” to the surface and the flexibility of the holoenzyme allows the PlyC EADs to cleave multiple bonds in a localized area weakening the overall superstructure of the PG. Alternatively, the enzymatic turnover could be dictated by a balance of on and off rates of the EADs and CBD monomers leading to widespread hydrolysis of the PG. Lastly, it is unknown whether PlyC binds, cleaves, and releases the PG at random points on the streptococcal surface or works its way down a single strand of PG in a processive manner. It is noteworthy that cellulase enzymes, which cleave the β1,4 glycosidic linkages in cellulose, possess a catalytic domain, a flexible linker, and a cellulose binding domain, analogous to the traditional endolysins. It has been demonstrated that energy is stored in the flexible linker can adopt compact and extended configurations that allows the cellulase to move in a “caterpillar-like” motion down a chain of cellulose [47, 48]. Although PlyC does not contain an equivalent flexible linker, the octameric nature of PlyCB invites the possibility that it may interact with the successive pRha strands allowing PlyC to depolymerize the PG in a zipper-like fashion.

In conclusion, the *α*1,2-1,3-pRha is the definite, minimal carbohydrate substrate for the PlyCB subunit. We validated this by comprehensive experiments, using pRha recombinantly produced in *E. coli*, and the *S. mutans* variants of the Rha-based polysaccharides. The work described here provides valuable insight into the molecular interactions that define a PlyC’s host specificity, which can inform the future studies as well as engineering approaches.

## Data Availability Statement

No mandated datasets are associated with the paper.

## ACKNOWLEDGEMENTS

We thank the CDC Streptococcus Lab and the Active Bacterial Core surveillance program for sharing isolates isolate 20170556 (stG6.0), 20171682 (stC74A.0), 20154376 (stG245.0), 20173686 (stG652.0), 20170560 and 20176966 (stG485.0). We also thank Ryan Heselpoth, Yang Shen, and Sara Linden for reagents, technical advice, and helpful discussion.

## Abbreviations (alphabetical order)

(CBD): Cell wall-binding domain
(EAD): enzymatically active domain
(Glc): Glucose
(GroP): glycerol phosphate
(GlyH): Glycosyl Hydrolase
(GAC): Group A Carbohydrate
(GAS): Group A Streptococcus
(GAVS): Group A-variant Streptococcus
(GCC): Group C Carbohydrate
(GCS): Group C Streptococcus
(GGC): Group G Carbohydrate
(GlcNAc): *N*-acetyl-glucosamine
(GalNAc): *N*-acetyl-galactosamine
(MurNAc): N-acetyl-muramic acid
(PG): Peptidoglycan
(pRha): polyrhamnose
(Rha): Rhamnose
(SDSE): *S. dysgalactiae subsp. equisimilis*
(SCC): *S. mutans* serotype c carbohydrate

