## Supplementary material for "Molecular basis for recognition of the Group A Carbohydrate backbone by the PlyC streptococcal bacteriophage endolysin": SF

A)

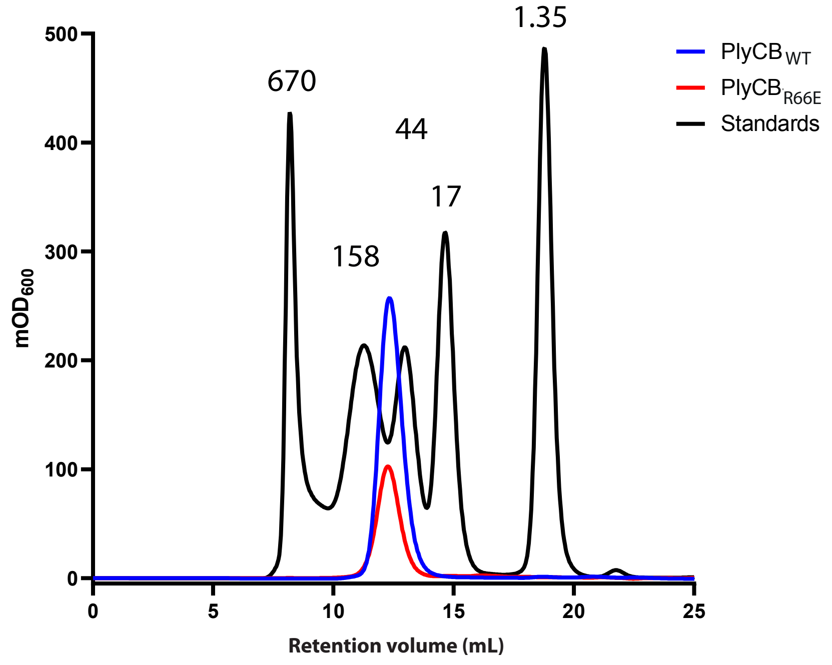


**SF 1) Gel filtration purification of WT and R66E-PlyCB proteins.** $PlyCB_{WT}$ (blue) and $PlyCB_{R66E}$ (red) proteins elute at a rentention volume of 12 mL. The PlyCB octamer has a theoretical mass of 63.8 kDa, with a monomeric mass of 7.9 kDa. Gel filtration standards are shown in kDa and include bovine thyroglobulin (670 kDa, void volume), bovine gamma-globulin (158 kDa), chicken ovalbumin (44 kDa), horse myoglobin (17 kDa), and vitamin B12 (1.35 kDa).

A)

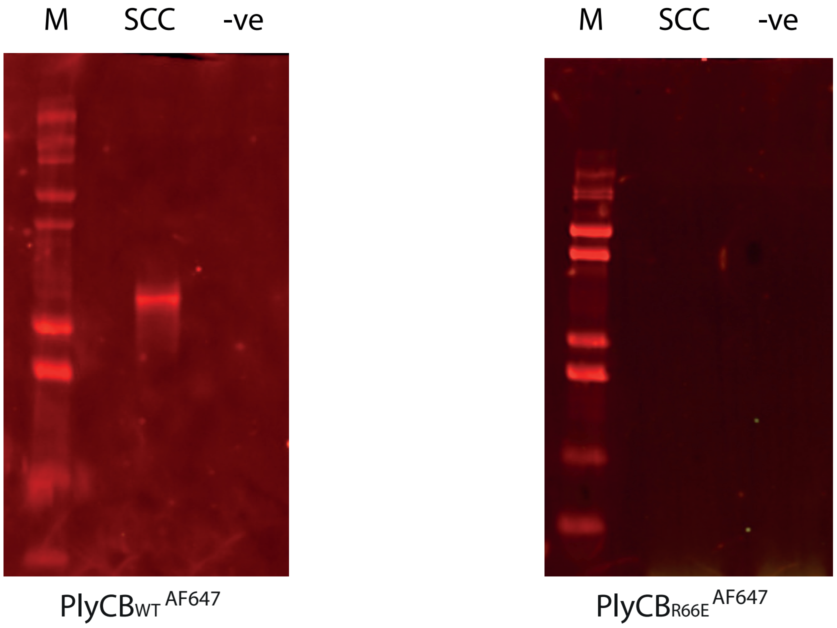


B)

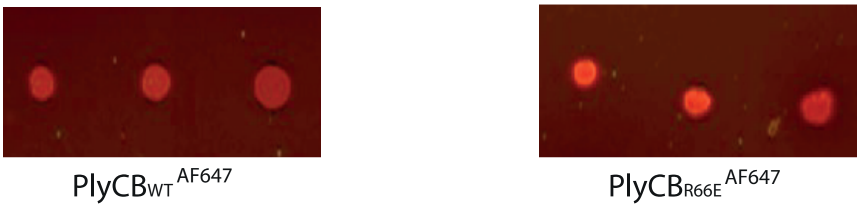


**SF 2)**

**PlyCB binds to pRha in *E. coli* cell lysate whilst $\text{PlyCB}_{\text{R66E}}$ does not. A)** Representative membrane blots showing that only PlyCB wildtype protein binds to total *E. coli* lysate when the pRha is expressed from the SCC gene cluster. $\text{PlyCB}_{\text{R66E}}$ fails to bind to pRha. **B)** Control dot blots showing that both PlyCB proteins are fluorescently labelled and detected when spotted directly onto nitrocellulose membrane (1:1000 dilution - identical to Figure A).

A)

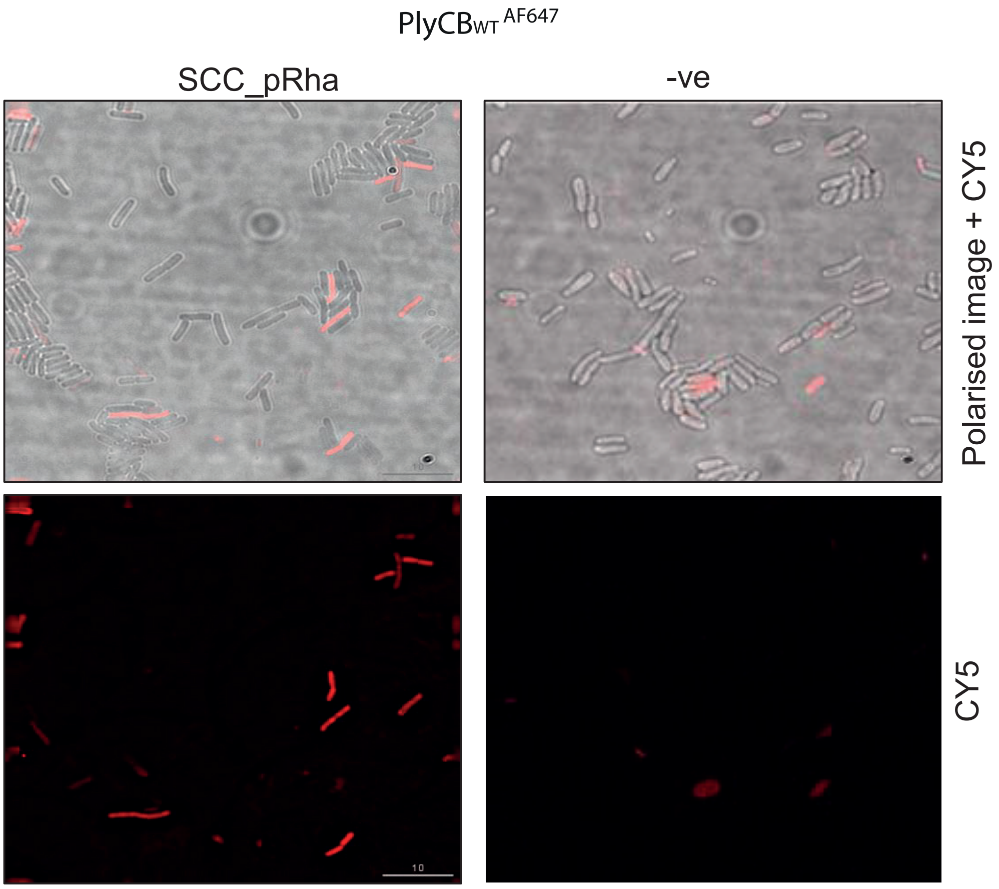


**SF 3)**

**PlyCB binds to pRha expressing *E. coli* cells analysed using microscopic analysis:** Representation of microscopic graph shows defined elongated $PlyCB_{WT}$ stained *E. coli* bacterial cells suggesting that PlyCB binds specifically to SCC_pRha cells but not to *E. coli* cells lacking pRha expression (-ve) detected using CY5 channel.
